# Polymicrobial-driven NLRP6 inflammasome regulates IL-1β production and alveolar bone loss in a murine model of periodontitis

**DOI:** 10.64898/2026.06.04.730168

**Authors:** Sarah Metcalfe, Rajendra P. Settem, Edwin Ovalle, Michelle Panasiewicz, Alejandro Escobar, Jason G. Kay

**Author notes:** Corresponding author: (JGK).

## Abstract

Periodontal disease is a chronic inflammatory condition that develops in response to oral microbiome dysbiosis and host-microbiome immune response dysregulation. The innate immune system plays a major role in the development and persistence of disease, in part by producing inflammatory cytokines. One of the major cytokines implicated in disease is interleukin-1β (IL-1β), which requires inflammasome activation. Much of the oral microbiome, including Streptococci, which are otherwise considered commensal, is required for the full development of periodontal disease. We have previously reported that inflammatory-activated macrophages and neutrophils counterintuitively allow survival of internalized *Streptococcus gordonii* over non- activated phagocytes. This internal bacterial survival leads to inflammasome activation via the cytoplasmic activator NLRP6, but not NLRP3, and subsequent increases in IL-1β release. Here, we test and find that the keystone pathogen *Porphyromonas gingivalis* can activate macrophages in a manner that allows for increased *S. gordonii* survival and IL-1β production above levels when *P. gingivalis* interacts with macrophages alone. We also use the mouse ligature-induced periodontal disease model to test the importance of NLRP6 in disease development. We found mice lacking NLRP6 had significantly reduced bone loss, IL-1β, and neutrophil infiltration following disease induced by *P. gingivalis* when *S. gordonii* or other mouse commensals were present, but had no effect when *S. gordonii* was inoculated alone. This work thus reveals an additional important inflammasome activation mechanism by which oral keystone pathogens may stimulate periodontal disease progression.

**Author Summary:** Chronic inflammation is a driver of many diseases, including periodontal disease. Periodontal disease is a long-lasting inflammatory disease caused by an unhealthy imbalance in the oral microbiome and an abnormal immune response toward those bacteria. A major inflammatory molecule involved in this inflammation is IL-1β, which is produced after inflammasome activation. We previously found that immune cells such as macrophages and neutrophils can unexpectedly allow *Streptococcus gordonii*, a normally health-associated oral bacterium, to survive within the immune cells and to trigger the NLRP6 inflammasome, leading to increased IL-1β release. In this study, we found that *Porphyromonas gingivalis*, a key periodontal pathogen, stimulates macrophages to allow *S. gordonii* survival and increased IL-1β production. Using a mouse model of periodontal disease, we also found that without NLRP6 mice had less bone loss, less inflammation, and fewer neutrophils when *P. gingivalis* was present along with other oral bacteria. These results show that NLRP6 plays an important role in how oral bacteria work together to worsen periodontal disease.

## Introduction

Periodontal disease is the most common chronic inflammatory condition not caused by a single organism, but through the development of a dysbiotic, or imbalanced, community of microorganisms. Dysbiotic oral microbiomes include some normally commensal organisms, along with keystone pathogens that are capable of driving this dysbiosis as minority members (1–5). *Porphyromonas gingivalis* is the most well-characterized keystone pathogen in periodontal disease development. *P. gingivalis* has the ability to inhibit some cellular signaling of, and survive within, immune cells such as dendritic cells and macrophages (6–12). This disruption of inflammatory response contributes to the emergence of bacterial dysbiosis and a resultant positive feedback loop of inflammation leading to more dysbiosis and ultimately the development of periodontal disease (2, 13, 14). Periodontal disease is not caused solely by keystone pathogens, but also involves opportunistic bacteria, including normally commensal species that contribute under dysbiotic conditions (2). Streptococci are part of the normal oral flora; however, they can also colonize extra-oral sites and contribute to systemic disease (15–19). *Streptococcus gordonii*, a commonly studied model oral streptococcus, can promote periodontitis by assisting *P. gingivalis* colonization and growth, acting as an accessory pathogen to enhance the pathogenicity of *P. gingivalis* (20–22). In addition, oral streptococci, while generally associated with the healthy oral microbiome due to them making up a large percentage of the microbiome, are present in higher absolute numbers during gingivitis and periodontal disease, even though they make up a lower percentage of the overall microbiome (5, 23–25).

Macrophages are an important cell type in the oral cavity, playing essential roles in maintaining mucosal immunity, including tolerance, and tissue homeostasis by physically interacting with and responding to oral microbes (26–30). In gingivitis and periodontal disease, the number of inflammatory or classically activated (M1) macrophages in the oral cavity increases, along with the inflammatory immune response they promote (31–34). In fact, depletion of macrophages reduces alveolar bone resorption by modulating the host immune response (35), and recruitment of non-activated or alternatively activated (M0 or M2) macrophages by CCL2 reduces alveolar bone loss in mouse models of periodontitis (36).

As part of the inflammatory response, macrophages produce cytokines and chemokines to orchestrate the immune response (30, 37). One such cytokine, IL-1β, is produced in an inflammasome-dependent manner and promotes inflammation, stimulates fever, and recruits and activates other immune cells (37, 38). Greatly increased IL-1β occurs in gingivitis and chronic periodontitis (33, 39, 40), and IL-1β has emerged as a possible therapeutic target in periodontal disease and other chronic inflammatory diseases (41–44). Contributing to periodontal disease-associated inflammation are infiltrating inflammatory macrophages, which have increased inflammasome components along with high levels of IL-1β production that lead to further promotion of inflammation and alveolar bone resorption (32, 33, 39, 41, 45).

We have found that *S. gordonii* is better able to survive within and damage the phagosomes of inflammatory macrophages over non-activated macrophages, due to differences in reactive oxygen species (ROS) production within the phagosomes (46). This increased survival and phagosomal damage lead to greater IL-1β production through activation of the inflammasome via the NLRP6 receptor in macrophages (47). NLRP6 is activated by LTA (Lipoteichoic acid), generally produced by Gram-positive bacteria (48). While NLRP6 is highly expressed in intestinal epithelial and goblet cells, where it is involved in intestinal microbial homeostasis, epithelial cell repair and regulation of mucus production (49–52), NLRP6 also functions within macrophages, where it can limit commensal driven inflammation while also having both protective and detrimental roles, as has been seen in *Listeria monocytogenes* infections (48, 53, 54). As our previous studies have examined activated macrophages’ responses solely to streptococci *in vitro*, here we begin to examine the importance of *P. gingivalis* co-incubation in altering NLRP6 responses to *S. gordonii* both *in vitro* and *in vivo*.

## Methods

### Cell Culture

RAW264.7 macrophages (ATCC) were grown in RPMI medium (Lonza or Corning) supplemented with 10% fetal bovine serum (FBS) (Corning) and 2 mM L-glutamine (Corning) at 37°C in 5% CO_2_. Prior to bacterial killing assays macrophages were stimulated with 20 ng/ml recombinant mouse IFN-γ (GenScript) for 24 hours and with 0.1 µg/ml lipopolysaccharide (LPS) (Salmonella enterica serotype Minnesota strain Re595; MilliporeSigma) for 2 hours to convert to activated (M1-like) macrophages.

Human monocytic THP-1 cells (ATCC) were maintained in RMPI medium supplemented with 10% FBS, 2 mM L-glutamine, 1 mM sodium pyruvate, 10 mM HEPES (Fisher Scientific), 1.5 g/L sodium bicarbonate and 0.05 mM 2-mercaptoethanol. Two days prior to an experiment, cells were differentiated to macrophages with 100 nM Phorbol 12-myristate 13-acetate (PMA) (Cayman Chemicals) for 24 hours then allowed to rest in media without PMA for an additional 24 hours (55). Cells were stimulated with 20 ng/ml human IFN-γ for 24 hours then with 0.1 µg/ml LPS for 2 hours to activate towards an M1-like macrophage or left unstimulated (56). For studies looking at cytokine production, cells were left unstimulated prior to the addition of bacteria.

Wild type (C57BL/6J) immortalized mouse bone marrow derived macrophages (iBMDMs) (provided by the lab of Dr. Gabriel Núñez, University of Michigan Medical School) were grown in RMPI medium supplemented with 10% FBS, 2 mM L-glutamine, 1 mM sodium pyruvate (Fisher Scientific). Cells were stimulated with 20 ng/mL mouse IFN-γ (GenScript) for 24 hours before experiments to differentiate to an M1-like macrophage or left unstimulated (M0 macrophages) (57).

### Microbial Culture

*S. gordonii* strain DL1 (provided by Dr. Stefan Ruhl, University at Buffalo) were grown in brain heart infusion (BHI) medium (BD Biosciences) supplemented with 0.5% yeast extract (MP Biomedicals) at 37°C and 5% CO_2_. *P. gingivalis* (ATCC 33277 or W50) were grown anaerobically at 37°C in tryptic soy broth supplemented with menadione (1 µg/ml) and hemin (10 µg/ml). All experiments used mid-log phase bacteria cultures and MOI was calculated by counting using Petroff-Hausser Counter.

### Bacterial Killing Assay

To determine bacterial survival within macrophages a modified gentamycin resistance assay was used (46, 58, 59). Briefly, macrophages were seeded in duplicate on 12-well plates. For activation (IFN-γ/LPS), macrophages were stimulated overnight with 20 ng/ml IFN-γ (human (BioLegend) or mouse (GenScript) as required) then with 0.1 µg/ml LPS for 2 hours. For experiments with dual incubation of *S. gordonii* and *P. gingivalis*, macrophages were not activated with IFN-γ or LPS. Mid-log growth *S. gordonii* was sonicated to break up chains and added to macrophages at a multiplicity of infection (MOI) of 10:1. Plates were centrifuged to synchronize contact of bacteria with macrophages (125 x g for 1 min). Cells were incubated at 37°C for 30 min, then one set of wells was washed extensively with PBS to remove external bacteria, macrophages were lysed with sterile H_2_O and serial diluted and plated on BHI or Todd-Hewitt (TH) plates to determine the initial number of bacteria taken up by the macrophages. For the other set of wells, 150 µg/ml gentamycin was added and incubated for 30 min at 37°C, after which the media was replace with fresh RPMI and incubated for an additional 1.5 hours. Again, macrophages were lysed with sterile H_2_O and serial diluted and plated on bacterial media plates and incubated overnight. After incubation, the number of CFU taken up and CFU survival 2 hours post-phagocytosis were determined. The ratio of surviving (2.5 hours) bacteria to phagocytosed (0.5 hours) bacteria gave us percent survival of bacteria within macrophages.

### Cytokine Analysis

To measure cytokine release from THP-1 macrophages, cells were seeded on 12 or 24-well plates at 5×10^5^ cells/ml and left unstimulated for 2 hours at 37°C. Bacteria were then added at an MOI = 10:1, plates were centrifuged to synchronize contact of bacteria with macrophages (125 x g for 1 min) then incubated at 37°C for 6, 12, or 24 hours. After incubation cell supernatants were collected and spun down to remove cell debris and bacteria. Levels of TNFα, IL-6 and IL-1β were measured by ELISA (R&D systems) according to the manufacturer’s instructions. Concentrations (pg/ml) were normalized to amount per 1x10^5^ macrophages.

### Flow Cytometry

Adherent cells were lifted by incubation with 0.5 mM EDTA at room temperature for 10 minutes and transferred to a 96-well v-bottom plate. All centrifugation was done at 4°C, 1500 rpm, for 8 minutes. Cells were blocked with mouse or human Fc Block (BD Biosciences) for 10 minutes at 4°C in 1% BSA in PBS. Cells were subsequently stained at 4°C and washed in BSA. For internal staining cells were fixed with 4% PFA for 20 minutes, then permeabilized with BD Perm/Wash buffer (BD Biosciences) according to manufacturer’s directions before staining with internal antibody. Flow cytometry was performed using a BD Fortessa flow cytometer, and all data were analyzed using FlowJo version 10.1 or higher.

### Ligature model of periodontal disease

Mouse experiments were approved by the University at Buffalo Institutional Animal Care and Use Committee (IACUC) (protocol ID: PROTO202100042). Wild type control C57BL/6J (Jackson labs strain #000664) and *Nlrp6-/-* mice (generously provided by Gabriel Núñez, University at Michigan [48]), 6-8 weeks in age, were divided into 5 groups (8-10 mice per group (4-5 female and 4-5 male)) as i) sham, ii) ligature alone, iii) ligature + *P. gingivalis,* iv) ligature + *S. gordonii*, and v) ligature + *P. gingivalis* & *S. gordonii*. After treatment with antibiotics to reduce, but not eliminate, the existing oral microflora (Kanamycin Sulphate 1 mg/ml for 5 days followed by a 3-day antibiotic free period), mice were intraperitoneally anesthetized with a 200 µl mixture of ketamine (10 mg/ml) and xylazine. A black braided silk ligature (6.0, Fisher Scientific) was placed on the left side second maxillary molars inoculated with 100 µl (1x10^9^ CFU/ml) of *S. gordonii* or *P. gingivalis,* or both, while the sham group received 2% CMC alone (Day-0). Booster doses with 100 µl of *P. gingivalis*, *S. gordonii* or both (1x10^9^ CFU/ml) were administered on the following Day-1 and 2 to the respective groups. After 14 days of ligature placement mice were sacrificed and the maxillary jaw bones were scanned with microCT. Briefly, the entire mouse head was harvested and fixed in 4% formaldehyde for high-energy microCT (Scanco100 μCT). Images were reconstructed and analyzed using Analyze Pro software (Analyze Direct, Inc. KS USA) to calculate the distance (micrometers) between the CEJ to ABC on the secondary molar at M-P (meso palatal), D-P (Disto-palatal), M-B (Meso-Buccal), D-P (Disto Buccal) sites. The total alveolar bone loss was calculated by adding the distances at meso and distal points on both palatal and buccal sides.

Bone volume calculations were performed on the microCT image stacks using FIJI (ImageJ) (60). Briefly, a reference-based registration (61) was applied to ensure consistent orientation and region-matched quantification across samples. A sham-control image was selected and manually oriented as the reference. A binary registration mask was created on the reference to outline the alveolar region surrounding the molars. Each sample was first manually aligned with the reference, then automatically refined using the registration mask, resulting in images with teeth and surrounding bone aligned. For quantification, a separate binary analysis mask was created on the reference image to define a standardized region of interest (ROI) around the second molar. The second molar was segmented and excluded before measurements. After registration, a FIJI script, available on <u>GitHub</u> was used to batch-apply the analysis mask to all samples and extract bone/tissue density and bone volume fraction (BV/TV) from the same region in each microCT image.

### Bulk RNA Preparation

RNA was extracted from collected tissue with Qiagen RNeasy kits. The extracted RNA was treated with DNAse1 using the New England Biolabs Monarch Spin RNA Cleanup Kit (T2020L). We used 100 ng of the total RNA as input for the Illumina Total Stranded RNA library prep kit with Ribo-Zero following the manufacturer’s protocol. Per-cycle basecall (BCL) files generated by the NovaSeq 6000 were converted to per-read FASTQ files using bcl2fastq version 2.20.0.422 using default parameters (Illumina). The quality of the sequencing was reviewed using FastQC version 0.11.9 (Babraham Bioinformatics). Detection of potential contamination was done using FastQ Screen version 0.14 (62). FastQC and FastQ Screen quality reports were summarized using MultiQC version 1.14 (63).

No adapter trimming was performed. Genomic alignments were performed using HISAT2 version 2.2.1 using default parameters (64). Ensembl reference GRCm38 was used for the reference genome and gene annotation set. Sequence alignments were compressed and sorted into binary alignment map (BAM) files using samtools version 1.6.1. Counting of mapped reads for genomic features was performed using Subread featureCounts version 2.0.4 (65) using the parameters -p -s 2 –g gene_name –t exon – Q 60 -B -C, the annotation file specified with –a was the Ensembl GRCm38 reference provided by Illumina’s iGenomes. Alignment statistics and feature assignment statistics were again summarized using MultiQC.

### Bulk RNA Sequencing Analysis

Samples were analyzed using R (V4.5.0) with DESeq2 package (1.48.2) (66). Basic quality control was performed by calculating total library size, the number of detected genes, and DESeq2 size factors for each sample. Sample counts were normalized using regularized log “rlog” and the within-group variability was calculated to perform differential expression analysis within each genotype. Differential expressions were determined by log fold changes, which were shrinkage-estimated using the apeglm method to improve the stability of effect size estimates (67). Genes were classified as significantly up- or downregulated based on an adjusted p-value threshold and a minimum fold-change threshold. Volcano plots were used to visualize the upregulated and downregulated genes using ggplot2 (v3.5.2) package. Gene Ontology (GO) of biological processes and KEGG enrichment analysis were performed between WT and *Nlrp6-/-* samples. To focus on immune-related biology, GO results were additionally filtered to the “immune system process” term. All enrichment plots were presented as bar charts using clusterProfiler (v4.16.0), with data obtained from the biomaRt database (68). We set significance thresholds for all analyses, and significantly enriched terms were defined by an alpha value (Benjamini-Hochberg correction) of less than 0.05.

### Tissue staining

Sections from fixed samples were stained with H&E using Hematoxylin (Gill III) (Sigma Aldrich) and Eosin-Y (Epredia), mounted using an immunohistochemistry mounting medium (Cell Signaling) and imaged on a Zeiss Axioskop. For Immunofluorescence staining, paraffin fixed decalcified slides were deparaffinized, then incubated in 0.2 M boric acid (Fisher Scientific) overnight at 60°C for antigen retrieval (69). Primary antibody IL-1β (clone 3A6, Cell Signaling), neutrophil elastase (clone E8U3X, Cell Signaling) or F4/80 (clone T45-2342, BD Pharmingen) were used at a dilution of 1:200 with 0.5% Bovine Serum Albumin (Fisher) overnight at 4°C in a moisturized chamber. Secondary antibody (Alexa-488 or Alexa-594 conjugated donkey anti-mouse, Alexa-488 donkey anti-rabbit or Alexa-647 donkey anti-rat (Jackson ImmunoResearch)) were added in a dilution of 1:500 in 0.5% Bovine Serum Albumin (Fisher) for 1 hour at room temperature. Slides were counterstained with DAPI (Vector Laboratories) and mounted with 50% glycerol mounting media containing N-propyl gallate (MP Biomedicals). Images were taken on an Andor Dragonfly Confocal Microscope at the Optical Imaging and Analysis Facility (School of Dental Medicine, University at Buffalo). Image analysis was performed using FIJI (60).

## Results

### *P. gingivalis* macrophage stimulation enhances survival of *S. gordonii*

Direct interaction of *S. gordonii* and *P. gingivalis* can result in periodontal disease due to the ability of *S. gordonii* to facilitate the colonization and pathogenicity of *P. gingivalis* (2, 21, 70, 71), though it is not clear if *P. gingivalis* can affect *S. gordonii* survival. Given our previous observations of increased *S. gordonii* survival and induction of IL-1β production in cytokine-induced inflammatory macrophages (46, 47), we wanted to test if *P. gingivalis*-activated macrophages led to similar changes in *S. gordonii* survival. We first examined how *P. gingivalis* affects macrophage activation phenotype. There was an increase in inflammatory macrophage (M1) markers CD80 and iNOS after stimulation with *P. gingivalis* for both 2 and 24 hours (Figure 1), agreeing with previous research showing *P. gingivalis* promoting an M1 phenotype while inhibiting an M2 phenotype (72–75). This is similar to what has been found with cytokine activation of macrophages, which leads to prolonged reactive oxygen production in macrophage phagosomes (76) and allows for increased *S. gordonii* survival at multiple MOIs (46, 47).

**Figure 1.**
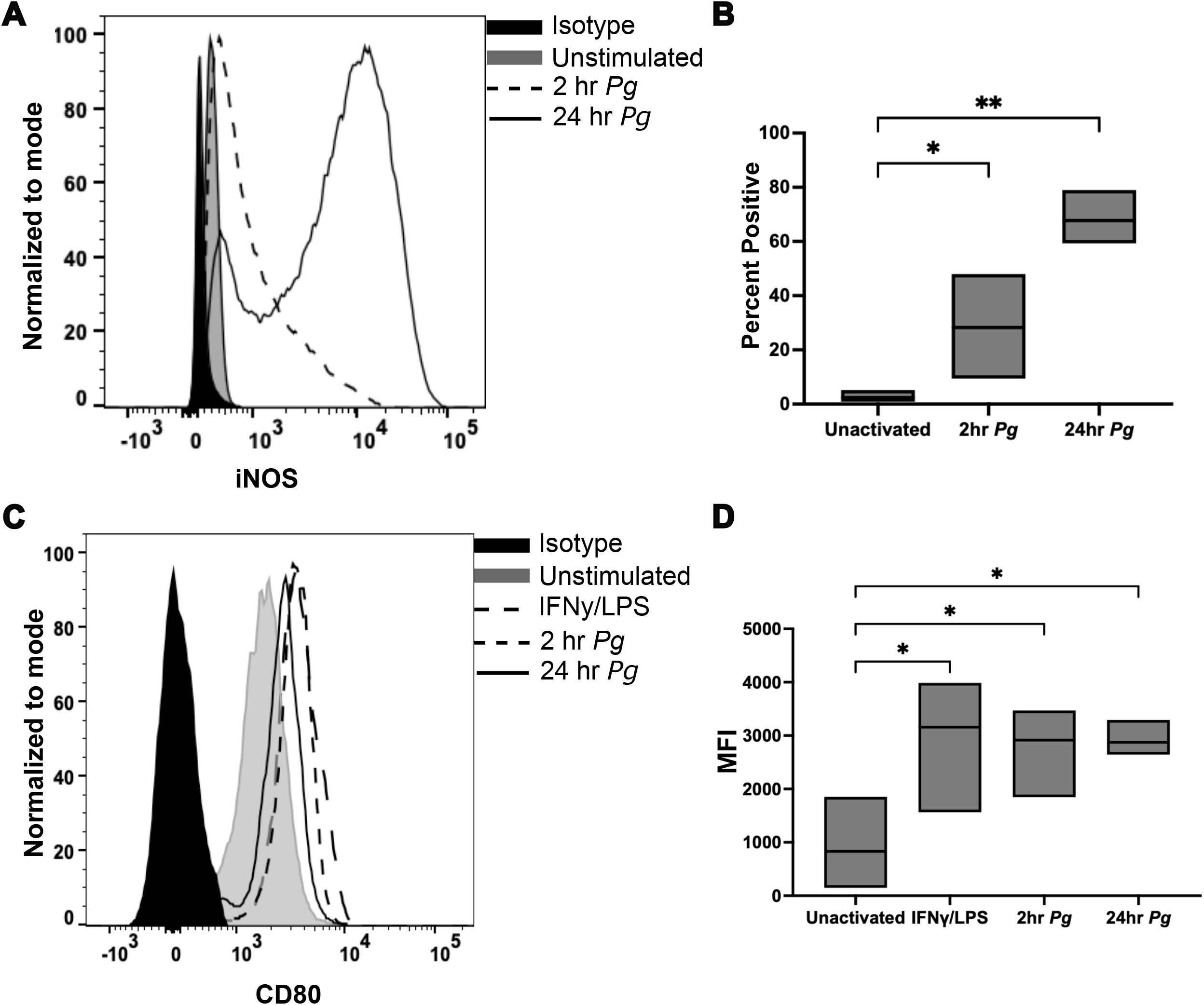
Inflammatory macrophage marker expression increases with *P. gingivalis* incubation. (**A-B**) Mouse bone marrow derived macrophages (BMDM) were differentiated with CSF-1, then stimulated with *P. gingivalis* for 2 or 24 hours. Expression of iNOS was observed by flow cytometry. Quantitation (B) showed a significant increase after 2 and 24 hours, with an average of 28% positive after 2 hours and 68% after 24 hours. (**C-D**) PMA differentiated THP-1 macrophages were stimulated with *P. gingivalis* for 2 or 24 hours, or IFNγ/LPS for 24 hours as a positive control, and the mean fluorescence intensity (MFI) of CD80 was determined by flow cytometry. Quantitation showed a significant increase in MFI over control after 2- and 24-hours incubation with *P. gingivalis*. A and C are representative plots, B and D are min to max boxes with means of 3 independent experiments. P values (*<0.05, **<0.01) were calculated by one-way ANOVA and Holm-Šídák’s multiple comparisons tests.

We next tested *S. gordonii’s* ability to survive within macrophages when co-cultured with *P. gingivalis* (Figure 2). For these experiments, two different combinations of *S. gordonii* and *P. gingivalis* were used: 80% *S. gordonii* with 20% *P. gingivalis*, or 50% of each bacterium, while maintaining an overall MOI of 10:1 (see methods). With co-incubation of *S. gordonii* and *P. gingivalis*, *S. gordonii* survival in both mouse and human macrophages increased after phagocytosis. Interestingly, at the closer-to-physiologically relevant ratio where *S. gordonii* outnumbers *P. gingivalis*, we saw the highest percent internal survival of *S. gordonii* (Figure 2A-C).

**Figure 2.**
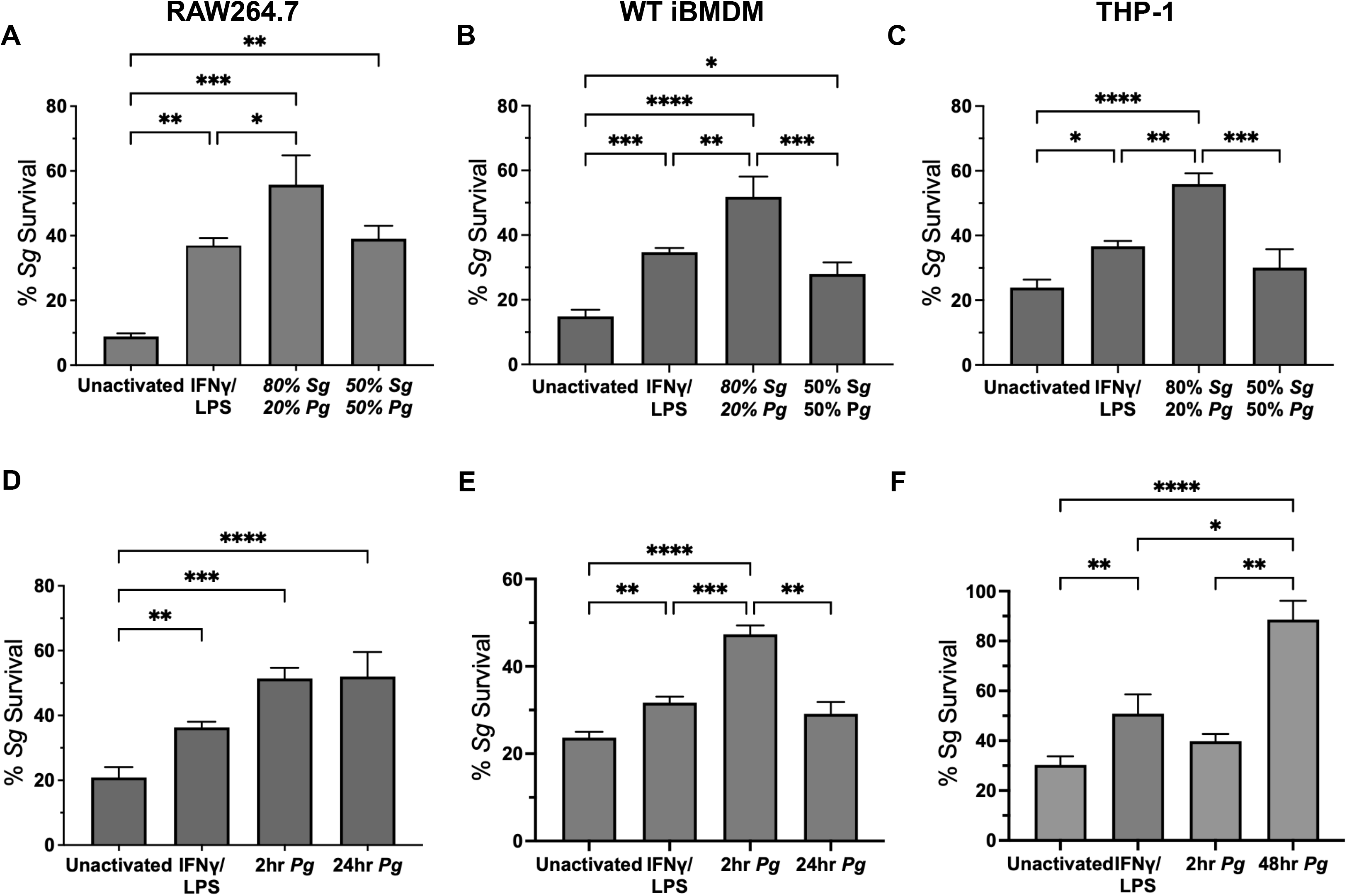
Survival of *S. gordonii* within macrophages increased with co-incubation of *P. gingivalis*. Shown is survival of *S. gordonii* within macrophages co-incubated with *P. gingivalis* at varying ratios (with a constant overall total MOI of 10:1) (A-C) or after stimulation with *P. gingivalis*, for the indicated times before incubation with *S. gordonii* (D-F); (**A,D**) RAW264.7 cells, (**B,E**) Immortalized mouse bone marrow derived macrophages (iBMDM) and (**C,F**) differentiated THP-1 cells. Shown are the means ± SEM of 3-8 independent experiments. P values (*<0.05, **<0.01, ***<0.001 and ****<0.0001) were calculated by one-way ANOVA and Tukey’s multiple comparisons tests.

We also tested *S. gordonii* survival in *P. gingivalis*-activated macrophages. Here we found that macrophages activated by *P. gingivalis* prior to the incubation of *S. gordonii* allowed for increased *S. gordonii* survival (Figure 2D-F), likely because *P. gingivalis* further stimulates macrophages to an inflammatory phenotype ((Figure 1) and (46, 47)).

### Inflammatory cytokine release increased upon co-incubation with *S. gordonii* and *P. gingivalis*

Activated macrophages release a myriad of pro-inflammatory cytokines to elicit an immune response and recruit other immune cells. We examined a panel of three pro-inflammatory cytokines prevalent in periodontal disease, IL-1β, TNFα, and IL-6 (33, 34, 77, 78) by human macrophages upon incubation with various combinations of *S. gordonii* and *P. gingivalis.* There were no significant changes in release of TNFα or IL-6 with co-incubation compared to incubation with *S. gordonii* or *P. gingivalis* alone (Figure 3A-B). There was, however, a significant increase in IL-1β production when *S. gordonii* was added simultaneously with *P. gingivalis* as compared to either bacterium alone (Figure 3C). An increase in IL-1β release was also seen when *P. gingivalis* strain W50 (Figure S1), a generally more pathogenic strain in lesions (79, 80). More significantly, when macrophages were stimulated with *P. gingivalis* for 24 hours prior to adding *S. gordonii,* there was a strong increase in IL-1β production (Figure 3D).

**Figure 3.**
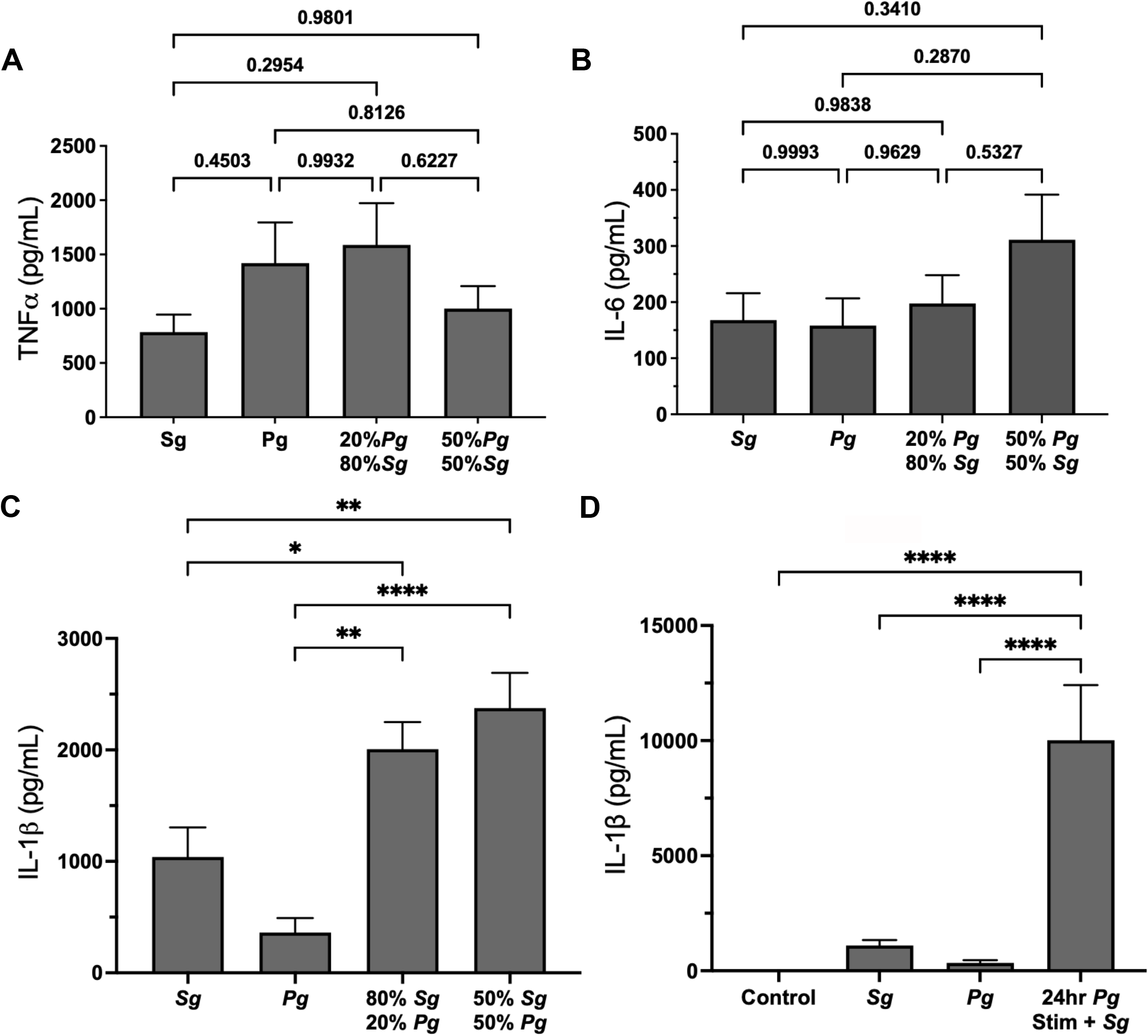
IL-1β, but not TNFα or IL-6, are increased following dual microbial incubation with macrophages. Inflammatory cytokine release from PMA differentiated THP-1 cells incubated with *S. gordonii, P. gingivalis,* or a combination thereof (All MOI = 10:1). (**A**) TNFα, (**B**) IL-6 and (**C**) IL-1β release after 6 hours incubation. (**D**) IL-1β release from THP-1 cells incubated with *S. gordonii* or *P. gingivalis* for 24 hours, or stimulated with *P. gingivalis* for 24 hours before incubating with *S. gordonii* for an additional 6 hours (24hr *Pg* Stim + *Sg*). Shown are means ± SEM of 4 or more independent experiments. P values (in A-B, or *<0.05, **<0.01, ***<0.001 and ****<0.0001 in C-D) were calculated by ordinary one-way ANOVA followed by Holm-Šídák’s multiple comparisons tests. No significant differences were seen with TNFα or IL-6 production.

### Co-incubation of *P. gingivalis* and *S. gordonii* in animal model of periodontitis

We have previously reported that *S. gordonii* survival is increased within inflammatory macrophages (46), whereupon it can damage the phagosome and stimulate the inflammasome for IL-1β release via the NLRP6 pathway (47). We found here that *P. gingivalis* can promote macrophages to an inflammatory phenotype (Figure 1), which is beneficial for *S. gordonii* survival and *S. gordonii*-induced IL-1β production (46, 47). We therefore next sought to determine whether disease development *in vivo* is dependent on NLRP6, specifically when both *P. gingivalis* and *S. gordonii* are present. Ligatures were placed on the second molar of either wild type (C57BL/6J) or NLRP6 knockout (*Nlrp6-/-*) mice and inoculated with a combination of *P. gingivalis* and *S. gordonii,* or each bacterium alone (see methods).

Following tissue collection, H&E staining and immunohistology for IL-1β were performed on gingival tissue sections from the mice (Figure 4A). Quantification of IL-1β fluorescence showed significantly reduced IL-1β in *Nlrp6-/-* mice that had ligatures with *P. gingivalis* as well as those that had ligatures with both *P. gingivalis* and *S. gordonii* (Figure 4B). Simultaneous immunohistology staining for IL-1β, F4/80 (to label monocyte/macrophages), and neutrophil elastase (to label neutrophils) of WT mice showed much of the IL-1β detected was within the F4/80 labelled macrophages (Figure 4C-D), confirming earlier reports (81, 82).

**Figure 4.**
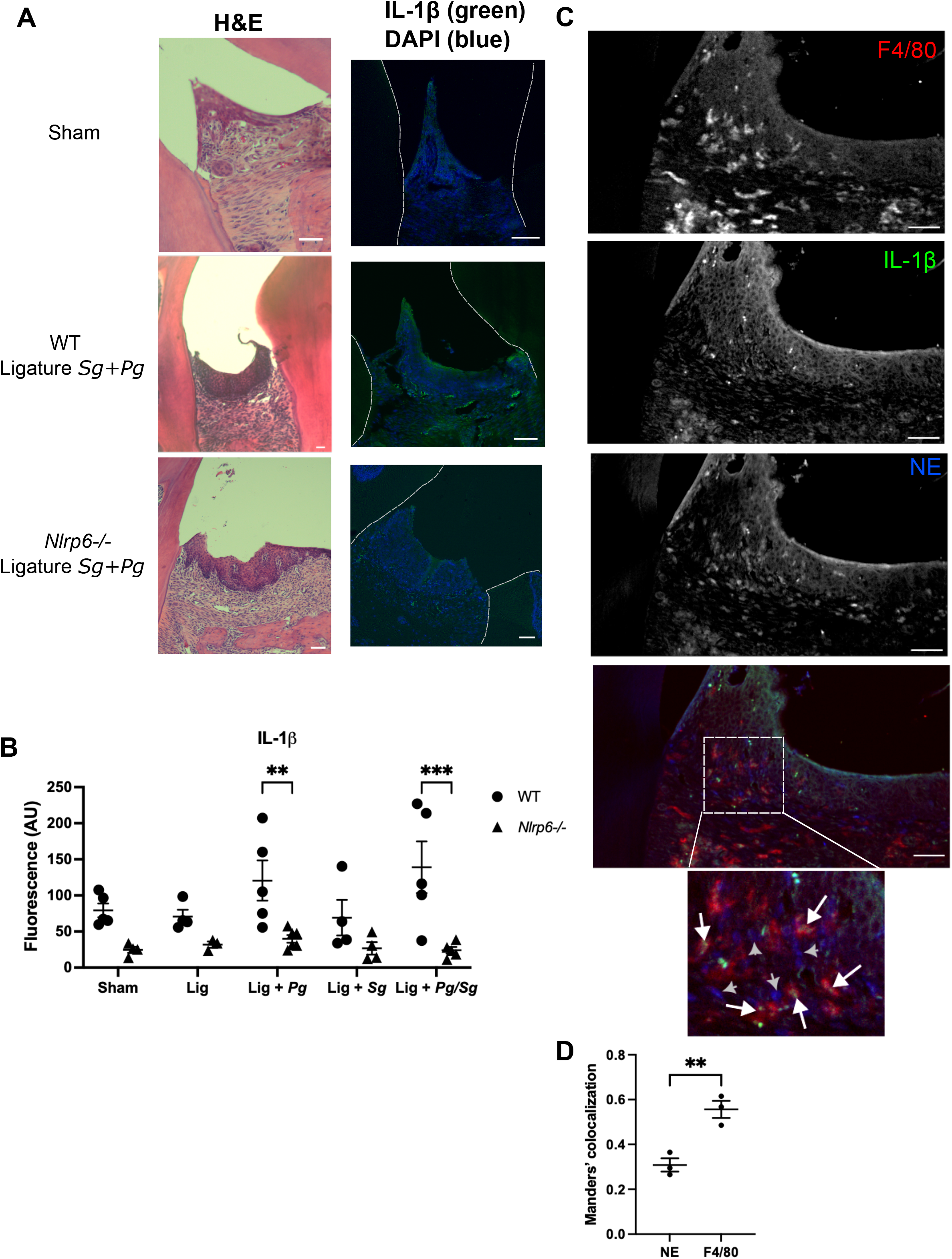
IL-1β production greatly reduced in gingiva of *Nlrp6-/-* mice with ligatures and *P. gingivalis*. (**A**) Representative images of H&E-stained sections and immunofluorescence of IL-1β (green) and nuclei (DAPI, blue). Bars = 100 μm. (**B**) Quantitation of IL-1β immunofluorescence within the gingiva. Shown are individual mice data with means ± SEM indicated. P values (*<0.05, **<0.01 and ***<0.001) calculated by ordinary two-way ANOVA followed by Holm-Šídák’s multiple comparisons test, with a single pooled variance. (**C**) Representative image of WT mice with ligatures showing immunofluorescence of F4/80 (red) to label monocytes/macrophages, neutrophil elastase (NE, blue) to label neutrophils and IL-1β (green). Inset shows F4/80 co-localizing with IL-1β (white arrows) while neutrophil elastase shows less overlap (gray arrowheads). Bars = 50 μm. (**D**) Manders’ colocalization of IL-1β immunofluorescence with neutrophils (NE) or macrophages (F4/80) confirms higher IL-1β colocalization with macrophages. P value (**<0.01) was calculated by ordinary unpaired t test.

MicroCT was also performed on the jaws to analyze bone loss (Figure 5). While all WT mice showed increased bone loss as measured by CEJ-ABC distance when a ligature was placed, mice that received both *P. gingivalis* and *S. gordonii,* but not *P. gingivalis* alone or *S. gordonii* alone, on the ligature had significantly increased CEJ-ABC distance than those with either ligature only (Figure 5B). *Nlrp6-/-* mice that had ligatures with *P. gingivalis,* or both *P. gingivalis* and *S. gordonii*, had significantly reduced CEJ-ABC distance than the equivalent wildtype mice (Figure 5B). Intriguingly, *Nlrp6-/-* mice with ligature alone or with *S. gordonii* only did not have decreased CEJ-ABC distance when compared to WT mice with the same treatment (p=0.9 and 0.07, respectively). Bone volume to tissue volume (BV/TV) analysis of the microCT images found that, similar to CEJ-ABC distance results, BV/TV ratios were significantly increased (indicating less bone loss) in *Nlrp6-/-* mice with ligatures incubated with *P. gingivalis* or both *P. gingivalis* and *S. gordonii* as compared to equivalent wildtype mice (Figure 5C). There were no significant changes in BV/TV in *Nlrp6-/-* mice in sham, ligature alone or ligature with *S. gordonii*, similar to the CEJ-ABC results.

**Figure 5.**
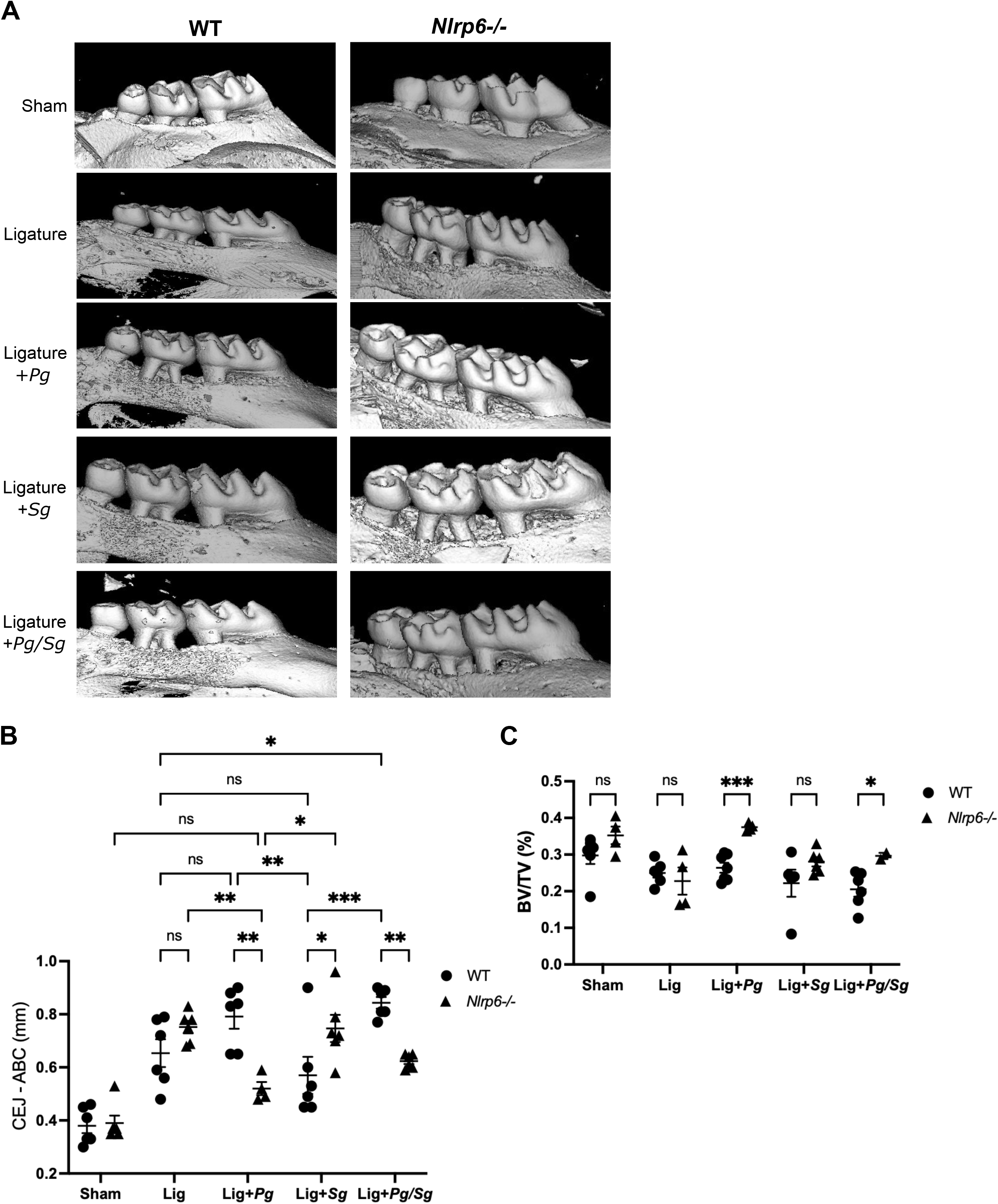
Bone loss in ligature-induced periodontal disease is significantly reduced in *Nlrp6-/-* mice. (A) Sample microCT images of WT and *Nlrp6-/-* maxilla after ligature (Lig) induced periodontitis. (B) Quantitation of cemental epithelial junction to alveolar bone crest (CEJ – ABC) distance measurements. Shown are means ± SEM, with P values (*<0.05, **<0.01 and ***<0.001; ns = not significant) calculated by ordinary two-way ANOVA followed by Holm-Šídák’s multiple comparisons test, with a single pooled variance. All mice with ligatures placed had significantly increased CEJ – ABC distances when compared to sham controls; for simplicity, only those not significant with sham are shown. (C) Quantitation of bone volume to total volume (BV/TV) fraction of microCT images. Shown are means ± SEM, with P values (*<0.05 and ***<0.001; ns = not significant) calculated by unpaired t test with Welch correction on each pair and Holm-Šídák’s multiple comparisons test.

We also examined bulk mRNA isolated from the gingival tissue of the mice (Figure 6). There were no significant differences in inflammatory cytokine production associated with periodontal disease (TNFα, IL-6, IL-17), with the exception of IL-1β which was reduced in *Nlrp6-/-* mice. There were also no significant changes in other inflammasome components, including ASC (pycard) or NLRP3. In general, all *Nlrp6-/-* samples exhibited more upregulated genes compared to their WT counterparts. Both *Nlrp6-/-* groups with *S. gordonii* (*Sg* and *Sg/Pg*) showed increased expression of genes related to cell motility, such as the Dnah family (Dnah1, Dnah3, Dnah5, Dnah6, Dnah7a, Dnah7b, Dnah11, Dnah12, Dnaic2, Drc1, Drc7) and the Cfap family (Cfap44, Cfap46, Cfap52, Cfap54, Cfap58, Cfap61, Cfap70), as well as genes involved in antimicrobial defense from the Bpifb family (Bpifb1, Bpifb9a, Bpifb9b) (Figure 6A,C). We conducted a GO enrichment analysis on the gene sets and the top pathways upregulated in the *Nlrp6-/- S. gordonii* samples were consistent with what was observed in the volcano plots, related to cell motility (Figure 6A,C). In contrast, the WT samples showed upregulation of pathways involved in fatty acid metabolism. When the GO analysis was further refined to focus on immune-related pathways, the most enriched pathways in *S. gordonii*-inoculated samples involved the classical inflammatory response (complement activation) and the alternative immune response (type II interferon, humoral immune response, and antibacterial humoral response) (Figure 6A,C).

**Figure 6.**
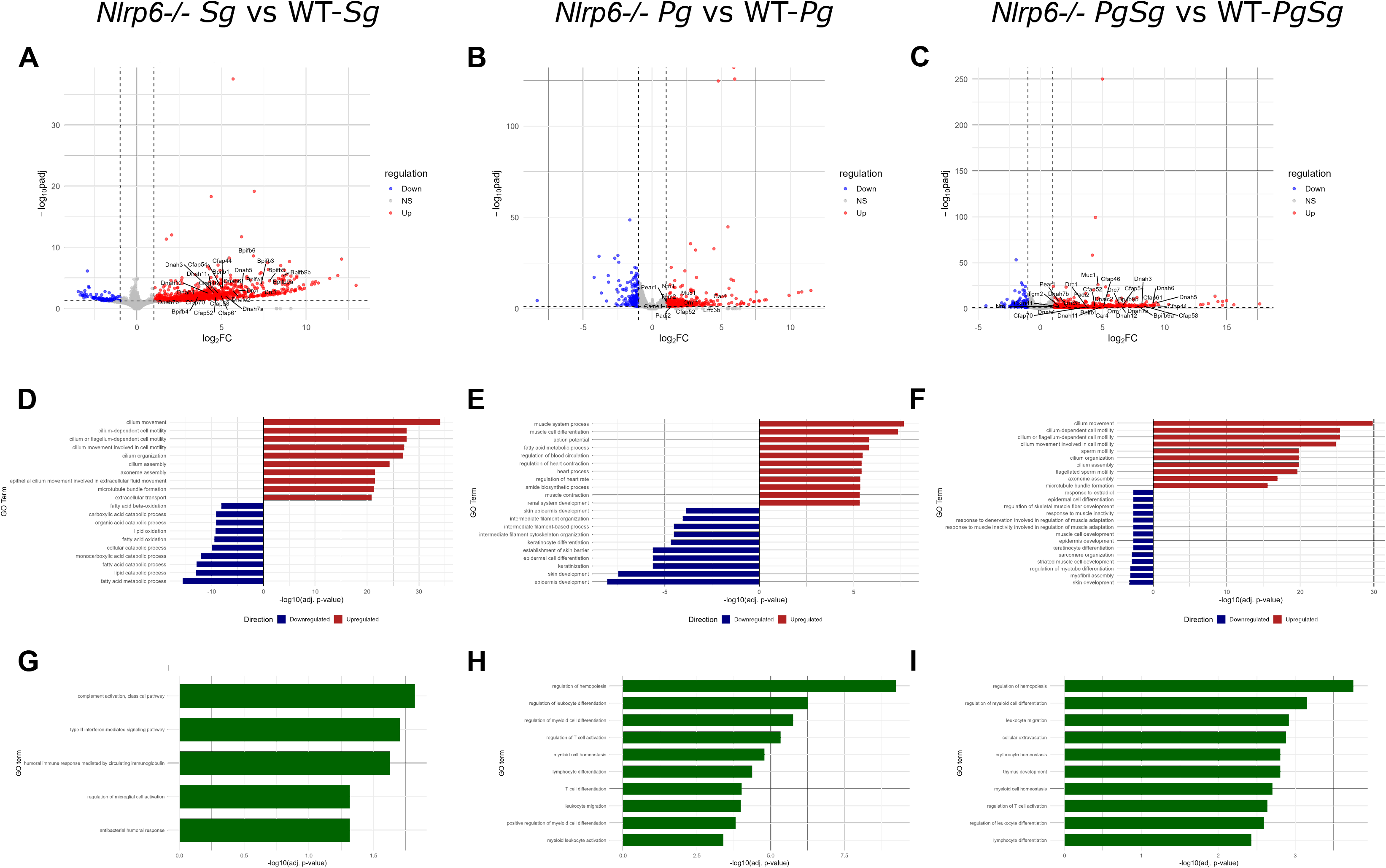
*Nlrp6-/-* mice alter immune responses during periodontal disease. (A-C) Volcano plots showing differential gene expression between *Nlrp6-/-* and WT treatments. Genes in blue are significantly downregulated; genes in red are significantly upregulated; genes in gray are not significantly changed between samples. (D-F) Gene Ontology (GO) enrichment analysis of biological processes was performed on *Nlrp6-/-* mice compared with WT mice. (G-I) GO enrichment pathways focused on the immune system process term. For all GO analyses, significantly enriched terms were defined by an adjusted P value (Benjamini-Hochberg correction) of less than 0.05.

The addition of *P. gingivalis* (*Pg* and *Sg/Pg*) induced inflammation, as seen with increased bone loss. Bulk-RNA analysis revealed that both *Nlrp6-/-* groups with *P. gingivalis* upregulated multiple genes involved in inflammation regulation (Csmd1, Padi2, Tgm2, and Orm1) and epithelial immune functions (Muc1, Pear1, Ntn1, Lrrc3b, and Car4) (Figure 6B,C). GO enrichment analysis revealed the top pathways upregulated in *P. gingivalis Nlrp6-/-* samples were related to metabolic changes and fatty acid metabolism (Figure 6B,C). WT samples had upregulated pathways related to epithelial function (Keratinocyte differentiation, establishment of skin barrier, epidermis development). *Pg-Sg Nlrp6-/-* samples upregulated pathways similar to *S. gordonii Nlrp6-/-* related to cilium movement, while the *Pg-Sg* WT samples had similar pathways upregulated to *P. gingivalis* WT samples in epithelial function (Figure 6B,C). The refined immunological GO pathway analysis revealed *P. gingivalis* samples showed an enrichment of pathways in the regulation of myeloid cell differentiation, leukocyte migration, and T cell signaling (Figure 6B,C).

Overall, the RNAseq data suggests that during *P. gingivalis* inflammation, in the *Nlrp6-/-* samples, the epithelial cells play a more crucial role, whereas in the WT sample, due to the excessive inflammation induced by IL-1β, the epithelial cells are attempting to restore the skin barrier. Additionally, although *S. gordonii* alone does not induce excessive inflammation and IL-1β production, *Nlrp6-/-* mice showed altered immune signaling, metabolism, and cellular motility in response to excessive Gram-positive bacteria.

Because we observed changes in cell motility pathways, we examined the levels of innate immune cell infiltration (neutrophils and monocytes/macrophages) using immunofluorescence on tissue sections with antibodies to neutrophil elastase and the macrophage marker F4/80 (Figure 7). Quantitation revealed a significant decrease in neutrophils within the marginal gingival tissue in *Nlrp6-/-* mice when a ligature was present as compared to WT mice (Figure 7B).

**Figure 7.**
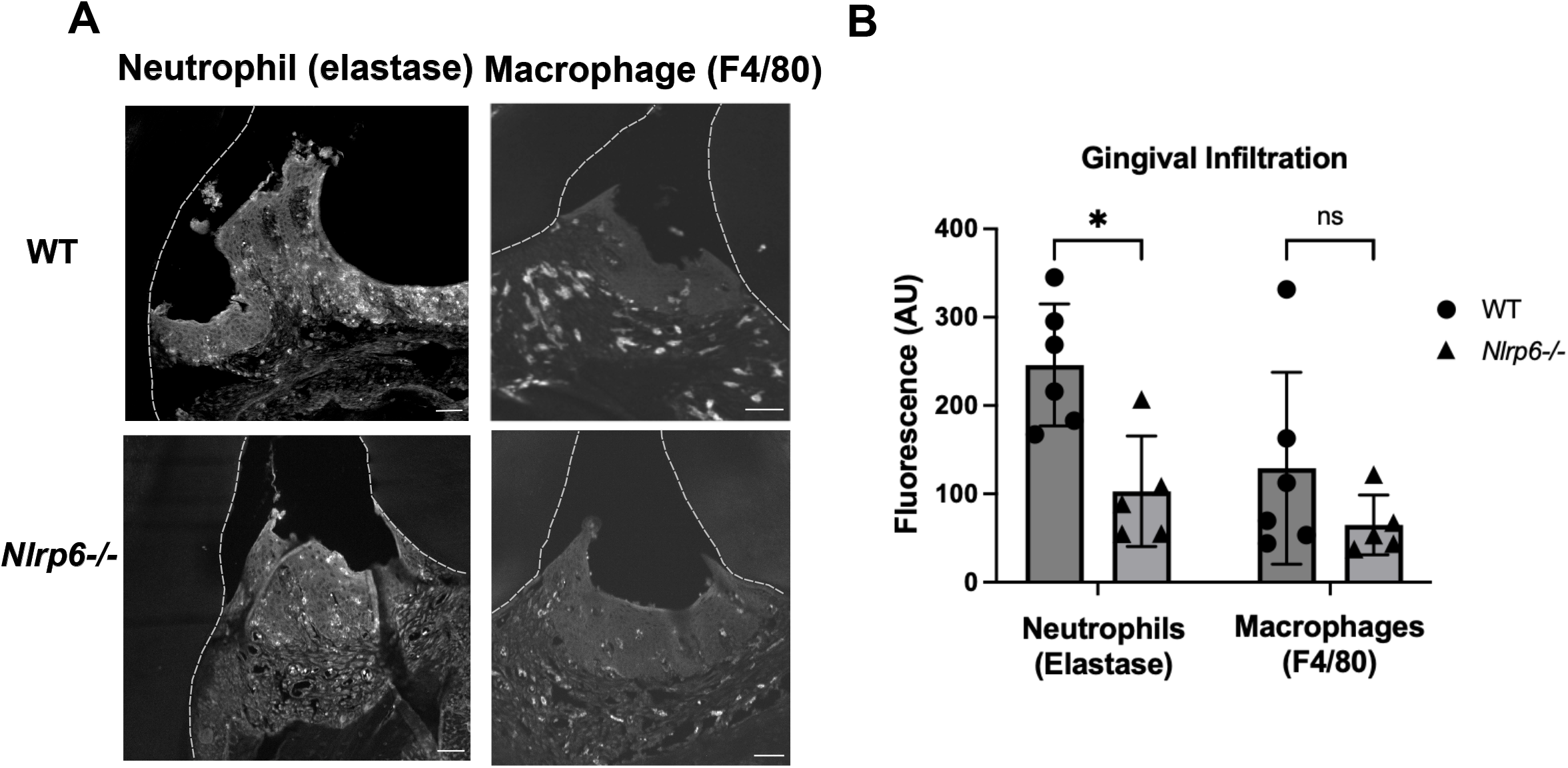
Gingiva neutrophil infiltration is reduced in *Nlrp6-/-* mice. (A) Representative images of mice with ligatures plus *S. gordonii* and *P. gingivalis* showing immunofluorescence of F4/80 to label monocytes/macrophages and neutrophil elastase to label neutrophils. Bars = 50 μm (B) Quantitation of F4/80 and neutrophil elastase immunofluorescence within the gingiva. Shown are means ± SEM, P values (*<0.05, ns = not significant) were calculated by t tests with Welch correction and Holm-Šídák’s multiple comparisons test.

## Discussion

IL-1β is well-known to be involved in the pathogenesis of chronic inflammatory diseases, including periodontal disease (41–44). While there are many activators of the inflammasome, the complex that activates caspases responsible for the maturation and release of IL-1β (83, 84), to date most studies investigating IL-1β and inflammasome involvement in periodontal disease have focused on the role of the activator NLRP3 (85–92). Our results reveal that the inflammasome activator NLRP6 is an important player in periodontal disease development.

In the oral cavity Gram-positive bacteria, which are the initial colonizers of the tooth surface, outnumber late colonizing Gram-negative anaerobes both in a healthy oral environment and during gingivitis development (93–95). In addition, while generally reduced as a percent of total bacteria present, Gram-positive bacteria are still a significant community member during advanced periodontal disease (2, 5, 94–96). Indeed, the numbers of all microbial members interacting with the immune system can be significant at all stages of health and disease (3, 13, 30), however, keystone pathogens such as *P. gingivalis* can dysregulate both the oral microbiome and the host immune response and promote disease (13, 97). Our initial *in vitro* results confirmed the ability of *P. gingivalis* to activate macrophages toward an M1 phenotype (Figure 1) and induce some IL-1β production (Figure 3). Similarly, *S. gordonii* incubation with macrophages alone, without pre-activation, was insufficient to elicit a robust IL-1β response (Figure 3 and (46)). We have also previously reported that changes in MOI of single bacterial incubations with macrophages, from 10 to 100, does not significantly change the levels of IL-1β release (98). In contrast, co-incubation with *P. gingivalis* and *S. gordonii* at a combined MOI of 10:1 induced substantially greater IL-1β production than either species alone, adding to our previous work showing that oral streptococci increase IL-1β production via activation of the NLRP6 inflammasome when engulfed by activated phagocytes (47, 99). In addition to an increase in IL-1β production, we observed an increase in *S. gordonii* survival in macrophages activated by *P. gingivalis* in a variety of macrophage cell lines, including the RAW264.7 cell line, which lacks ASC (100), a crucial central adaptor molecule for the inflammasome pathway, as we’ve seen previously (46, 47), and suggests *S. gordonii* survival is not dependent on inflammasome activation by *P. gingivalis*. Here, we saw macrophage activation by *P. gingivalis*, together with *S. gordonii*-mediated NLRP6 signaling, induced robust IL-1β release above what either bacterium was able to induce independently. Given that *P. gingivalis* is known to activate AIM2 and NLRP3-dependent inflammasomes (101–105), the enhanced IL-1β response may be due to the activation of multiple inflammasome pathways. These results are consistent with previous studies demonstrating enhanced cytokine production following exposure to multiple oral bacterial species (106, 107).

To understand the *in vivo* contribution of NLRP6 signaling to the inflammatory response, and periodontal disease development, induced by co-infection with *P. gingivalis* and *S. gordonii*, we used an NLRP6 knockout mouse model. In the presence of *P. gingivalis*, *Nlrp6-/-* mice exhibited significantly reduced disease severity, as measured by reduced bone loss and IL-1β production in the ligature model. These findings provide further evidence that *P. gingivalis* relies on the normally commensal oral microbiome (2), including oral streptococci, not only for colonization (21, 22), but also to promote dysbiosis. Our findings further demonstrated that the addition of *P. gingivalis* alone to ligatures did not significantly increase bone loss when compared to ligatures alone, though it did increase bone loss compared to ligatures with *S. gordonii* alone (Figure 5). The addition of ligatures can cause significant disease and bone loss due to increased microbial load (108–110), but without a robust microbiome to assist with its colonization, *P. gingivalis* cannot cause disease (2, 21, 111). Most mouse studies examining *P. gingivalis*-induced periodontal disease use a gavage model (without ligatures) over a time course of ∼6 weeks (2, 21, 89, 112, 113). The results presented here suggest cooperation between *S. gordonii* and *P. gingivalis* can cause increased disease over ligatures alone. Further, when combined with our previous results showing incubation with inflammatory-activated macrophages allows for increased *S. gordonii* survival within, and escape from, these macrophage phagosomes (46) and subsequent NLRP6 inflammasome activation (47), the results presented here suggest that the keystone bacterium *P. gingivalis* not only enhances inflammation in a positive feedback loop which further promotes dysbiotic microbial growth (13), but also allows *S. gordonii* to actively promote inflammation via NLRP6 inflammasome activation.

While periodontal bone loss was decreased in *Nlrp6^-/-^*mice as compared to WT mice when *P. gingivalis* was added to the ligatures, either alone or with *S. gordonii, S. gordonii* alone in *Nlrp6-/-* mice did not show bone loss changes when compared to WT mice. *S. gordonii* is part of the normal healthy microflora, and under these conditions interactions of the eubiotic microbiome with the immune system is generally homeostatic (14). As we have previously shown, NLRP6-dependent, *S. gordonii* mediated increases in IL-1β release from innate immune cells only occurs when the immune cells have been previously activated (46, 47, 99). This suggests that detrimental effects of NLRP6 in the oral environment may become important only during persistent inflammation. This aligns with previous research on the role of NLRP6 in the gut, which found it to have a regulatory and protective role in many infections but can be detrimental in cases of existing chronic inflammation (48, 49, 53, 114, 115). The requirement for enhanced inflammation *in vivo* likely stems from the presence of *P. gingivalis*, which is known to influence the immune system (1, 14, 116) and, as we showed here, can influence IL-1β production by macrophages in response to S*. gordonii*. When *P. gingivalis* is added alone without *S. gordonii*, the mouse’s naturally present commensals can partially enhance the effect of *P. gingivalis* (2). Indeed, we have seen that activated macrophages have altered responses to other oral

Streptococcal species (47), though a thorough investigation into the survival and IL-1β induction ability of additional oral Streptococci species is not yet complete. Similarly, the ability of Gram-negative keystone oral pathogens, other than *P. gingivalis,* to activate immune cells and allow for increased survival of *S. gordonii* (which is predominantly in gingival plaque (117)), or other oral Streptococci, leading to NLRP6-dependent IL-1β production is under active investigation.

Part of the difference in response with and without *P. gingivalis* added to the ligatures may also be due to a shift in the major cell type responding with NLRP6 signaling. Studies of gut infections, where NLRP6 is highly expressed in epithelial cells (118), suggest NLRP6 plays a mostly protective role (48, 53), while NLRP6 appears to promote disease when myeloid cell NLRP6 responses dominate (49, 115, 119). The relative expression of NLRP6 in different cells within the oral cavity is unclear, though it can activate the inflammasome and regulate cytokine responses in periodontal ligament cells and gingival fibroblasts (120, 121). Studies to understand NLRP6 within the oral cavity environment and its role in regulating the oral microbiome are therefore an important area for future research. Related, the interconnection between NLRP6 and other inflammasome activators implicated in periodontal disease, including NLRP3 and AIM2 that can be activated by *P. gingivalis* (87, 90, 101, 104, 105), and their relative importance in disease, is an area that needs clarification. Other than IL-1β, our results indicated other inflammatory cytokines often associated with periodontal disease, TNFα and IL-6, had no change in expression. This may partly explain why no changes were seen in *Nlrp6-/-* mice with ligature alone or with ligature and *S. gordonii* alone; these cytokines also play a role in periodontal bone loss (30, 82) and likely are important in the milder bone loss seen under these conditions.

Neutrophils are critical mediators of inflammation during periodontal disease development, and the number of neutrophils in *Nlrp6-/-* mice was significantly reduced, though not eliminated. This is important as increased neutrophils correlate with disease models and clinical data (122, 123), whereas neutropenia is also associated with poor oral health (124). However, how reduced IL-1β production in *Nlrp6-/-* mice, or the absence of NLRP6 itself, affects the adaptive immune cells involved in periodontal disease, such as Th17 cells, remains an open question. While IL-1β appears not to be important for the direct recruitment and expansion of Th17 cells, which are implicated in periodontal disease (125), we observed broad overall changes in T cell signaling in our RNAseq data. Parsing out how these changes specifically alter the adaptive immune landscape (30) will be important to understand in future studies. It will also be important to further understand the role of the NLRP6 inflammasome in periodontal disease-linked development of innate trained immunity and systemic diseases (126, 127).

In summary, our results indicate that NLRP6 plays an important role in IL-1β production and periodontal disease progression in the ligature-induced model. Since NLRP6 is activated by Gram-positive LTA (48), its role may lie in enabling normally commensal members of the oral microbiome to become pathobionts when a keystone pathogen, or other initiator of disease or inflammation, is present.

## Acknowledgements

Microscopy, microCT and flow cytometry were performed in the Optical Imaging and Analysis Facility in the School of Dental Medicine at the University at Buffalo. We also thank the director of the facility, Dr. Andrew McCall, for assistance with development of the bone volume calculation script. RNA isolation and sequencing was performed at the University at Buffalo Genomics and Bioinformatics facility.

---

**Figure S1. IL-1β release from macrophages increased with co-incubation of *P. gingivalis* strain W50.** IL-1β release from PMA differentiated THP-1 cells incubated with *S. gordonii*, *P. gingivalis* strain W50, or a combination thereof (all total MOI = 10:1) after 6 hours incubation. Shown are means ±SEM of 3 independent experiments. P values (*<0.05) were calculated by ordinary one-way ANOVA followed by Holm-Šídák’s multiple comparisons tests.

## References

1. Hajishengallis G, Lamont RJ. Dancing with the Stars: How Choreographed Bacterial Interactions Dictate Nososymbiocity and Give Rise to Keystone Pathogens, Accessory Pathogens, and Pathobionts. Trends Microbiol. 2016;24(6):477–89.

2. Hajishengallis G, Liang S, Payne MA, Hashim A, Jotwani R, Eskan MA, et al. Low-abundance biofilm species orchestrates inflammatory periodontal disease through the commensal microbiota and complement. Cell Host Microbe. 2011;10(5):497–506.

3. Ebersole JL, Dawson D, 3rd, Emecen-Huja P, Nagarajan R, Howard K, Grady ME, et al. The periodontal war: microbes and immunity. Periodontol 2000. 2017;75(1):52–115.

4. Irie K, Novince CM, Darveau RP. Impact of the Oral Commensal Flora on Alveolar Bone Homeostasis. J Dent Res. 2014;93(8):801–6.

5. Abusleme L, Hoare A, Hong BY, Diaz PI. Microbial signatures of health, gingivitis, and periodontitis. Periodontol 2000. 2021;86(1):57–78.

6. Wang M, Krauss JL, Domon H, Hosur KB, Liang S, Magotti P, et al. Microbial hijacking of complement-toll-like receptor crosstalk. Sci Signal. 2010;3(109):ra11.

7. Wang M, Shakhatreh MA, James D, Liang S, Nishiyama S, Yoshimura F, et al. Fimbrial proteins of porphyromonas gingivalis mediate in vivo virulence and exploit TLR2 and complement receptor 3 to persist in macrophages. J Immunol. 2007;179(4):2349–58.

8. Darveau RP, Belton CM, Reife RA, Lamont RJ. Local chemokine paralysis, a novel pathogenic mechanism for Porphyromonas gingivalis. Infect Immun. 1998;66(4):1660–5.

9. Widziolek M, Mieszkowska A, Marcinkowska M, Salamaga B, Folkert J, Rakus K, et al. Gingipains protect Porphyromonas gingivalis from macrophage-mediated phagocytic clearance. PLoS Pathog. 2025;21(1):e1012821.

10. Abdi K, Chen T, Klein BA, Tai AK, Coursen J, Liu X, et al. Mechanisms by which Porphyromonas gingivalis evades innate immunity. PLoS One. 2017;12(8):e0182164.

11. Zheng Y, Wang Z, Weng Y, Sitosari H, He Y, Zhang X, et al. Gingipain regulates isoform switches of PD-L1 in macrophages infected with Porphyromonas gingivalis. Sci Rep. 2025;15(1):10462.

12. Maekawa T, Krauss JL, Abe T, Jotwani R, Triantafilou M, Triantafilou K, et al. Porphyromonas gingivalis manipulates complement and TLR signaling to uncouple bacterial clearance from inflammation and promote dysbiosis. Cell Host Microbe. 2014;15(6):768–78.

13. Hajishengallis G, Lamont RJ, Koo H. Oral polymicrobial communities: Assembly, function, and impact on diseases. Cell Host Microbe. 2023;31(4):528–38.

14. Lamont RJ, Koo H, Hajishengallis G. The oral microbiota: dynamic communities and host interactions. Nat Rev Microbiol. 2018;16(12):745–59.

15. Han YW, Wang X. Mobile microbiome: oral bacteria in extra-oral infections and inflammation. J Dent Res. 2013;92(6):485–91.

16. Offenbacher S, Beck JD, Moss K, Mendoza L, Paquette DW, Barrow DA, et al. Results from the Periodontitis and Vascular Events (PAVE) Study: a pilot multicentered, randomized, controlled trial to study effects of periodontal therapy in a secondary prevention model of cardiovascular disease. J Periodontol. 2009;80(2):190–201.

17. Douglas CW, Heath J, Hampton KK, Preston FE. Identity of viridans streptococci isolated from cases of infective endocarditis. J Med Microbiol. 1993;39(3):179–82.

18. Baddour LM, Tayidi MM, Walker E, McDevitt D, Foster TJ. Virulence of coagulase-deficient mutants of Staphylococcus aureus in experimental endocarditis. J Med Microbiol. 1994;41(4):259–63.

19. Yombi J, Belkhir L, Jonckheere S, Wilmes D, Cornu O, Vandercam B, et al. Streptococcus gordonii septic arthritis: two cases and review of literature. BMC Infect Dis. 2012;12:215.

20. Hajishengallis G. Immunomicrobial pathogenesis of periodontitis: keystones, pathobionts, and host response. Trends Immunol. 2014;35(1):3–11.

21. Daep CA, Novak EA, Lamont RJ, Demuth DR. Structural dissection and in vivo effectiveness of a peptide inhibitor of Porphyromonas gingivalis adherence to Streptococcus gordonii. Infect Immun. 2011;79(1):67–74.

22. Kuboniwa M, Houser JR, Hendrickson EL, Wang Q, Alghamdi SA, Sakanaka A, et al. Metabolic crosstalk regulates Porphyromonas gingivalis colonization and virulence during oral polymicrobial infection. Nat Microbiol. 2017;2(11):1493–9.

23. Hong BY, Furtado Araujo MV, Strausbaugh LD, Terzi E, Ioannidou E, Diaz PI. Microbiome profiles in periodontitis in relation to host and disease characteristics. PLoS One. 2015;10(5):e0127077.

24. Abusleme L, Dupuy AK, Dutzan N, Silva N, Burleson JA, Strausbaugh LD, et al. The subgingival microbiome in health and periodontitis and its relationship with community biomass and inflammation. ISME J. 2013;7(5):1016–25.

25. Curtis MA, Diaz PI, Van Dyke TE. The role of the microbiota in periodontal disease. Periodontol 2000. 2020;83(1):14–25.

26. Dixon DR, Bainbridge BW, Darveau RP. Modulation of the innate immune response within the periodontium. Periodontol 2000. 2004;35:53–74.

27. Moutsopoulos NM, Konkel JE. Tissue-Specific Immunity at the Oral Mucosal Barrier. Trends Immunol. 2018;39(4):276–87.

28. Dutzan N, Konkel JE, Greenwell-Wild T, Moutsopoulos NM. Characterization of the human immune cell network at the gingival barrier. Mucosal Immunol. 2016;9(5):1163–72.

29. Merry R, Belfield L, McArdle P, McLennan A, Crean S, Foey A. Oral health and pathology: a macrophage account. Br J Oral Maxillofac Surg. 2012;50(1):2–7.

30. Baima G, Arce M, Romandini M, Van Dyke T. Inflammatory and Immunological Basis of Periodontal Diseases. J Periodontal Res. 2025.

31. Gonzalez OA, Novak MJ, Kirakodu S, Stromberg A, Nagarajan R, Huang CB, et al. Differential Gene Expression Profiles Reflecting Macrophage Polarization in Aging and Periodontitis Gingival Tissues. Immunol Invest. 2015;44(7):643–64.

32. Topoll HH, Zwadlo G, Lange DE, Sorg C. Phenotypic dynamics of macrophage subpopulations during human experimental gingivitis. J Periodontal Res. 1989;24(2):106–12.

33. Yang J, Zhu Y, Duan D, Wang P, Xin Y, Bai L, et al. Enhanced activity of macrophage M1/M2 phenotypes in periodontitis. Arch Oral Biol. 2018;96:234–42.

34. Yu T, Zhao L, Huang X, Ma C, Wang Y, Zhang J, et al. Enhanced Activity of the Macrophage M1/M2 Phenotypes and Phenotypic Switch to M1 in Periodontal Infection. J Periodontol. 2016;87(9):1092–102.

35. Lam RS, O’Brien-Simpson NM, Lenzo JC, Holden JA, Brammar GC, Walsh KA, et al. Macrophage depletion abates Porphyromonas gingivalis-induced alveolar bone resorption in mice. J Immunol. 2014;193(5):2349–62.

36. Zhuang Z, Yoshizawa-Smith S, Glowacki A, Maltos K, Pacheco C, Shehabeldin M, et al. Induction of M2 Macrophages Prevents Bone Loss in Murine Periodontitis Models. J Dent Res. 2019;98(2):200–8.

37. Arango Duque G, Descoteaux A. Macrophage cytokines: involvement in immunity and infectious diseases. Front Immunol. 2014;5(11):491.

38. Delaleu N, Bickel M. Interleukin-1 beta and interleukin-18: regulation and activity in local inflammation. Periodontol 2000. 2004;35:42–52.

39. Kinane DF, Winstanley FP, Adonogianaki E, Moughal NA. Bioassay of interleukin 1 (IL-1) in human gingival crevicular fluid during experimental gingivitis. Arch Oral Biol. 1992;37(2):153–6.

40. Zhong Y, Slade GD, Beck JD, Offenbacher S. Gingival crevicular fluid interleukin-1beta, prostaglandin E2 and periodontal status in a community population. J Clin Periodontol. 2007;34(4):285–93.

41. Hasturk H, Kantarci A, Goguet-Surmenian E, Blackwood A, Andry C, Serhan CN, et al. Resolvin E1 regulates inflammation at the cellular and tissue level and restores tissue homeostasis in vivo. J Immunol. 2007;179(10):7021–9.

42. Van Dyke TE. The management of inflammation in periodontal disease. J Periodontol. 2008;79(8 Suppl):1601–8.

43. Dinarello CA. Interleukin-1 in the pathogenesis and treatment of inflammatory diseases. Blood. 2011;117(14):3720–32.

44. Marchesan JT. Inflammasomes as contributors to periodontal disease. J Periodontol. 2020;91 Suppl 1(Suppl 1):S6–S11.

45. Lam RS, O’Brien-Simpson NM, Hamilton JA, Lenzo JC, Holden JA, Brammar GC, et al. GM-CSF and uPA are required for Porphyromonas gingivalis-induced alveolar bone loss in a mouse periodontitis model. Immunol Cell Biol. 2015;93(8):705–15.

46. Croft AJ, Metcalfe S, Honma K, Kay JG. Macrophage Polarization Alters Postphagocytosis Survivability of the Commensal Streptococcus gordonii. Infect Immun. 2018;86(3):e00858–17.

47. Metcalfe S, Panasiewicz M, Kay JG. Inflammatory macrophages exploited by oral streptococcus increase IL-1B release via NLRP6 inflammasome. J Leukoc Biol. 2023;114(4):347–57.

48. Hara H, Seregin SS, Yang D, Fukase K, Chamaillard M, Alnemri ES, et al. The NLRP6 Inflammasome Recognizes Lipoteichoic Acid and Regulates Gram-Positive Pathogen Infection. Cell. 2018;175(6):1651–64 e14.

49. Venuprasad K, Theiss AL. NLRP6 in host defense and intestinal inflammation. Cell Reports. 2021;35(4).

50. Fan W, Ding C, Liu S, Gao X, Shen X, De Boevre M, et al. Estrogen receptor β activation inhibits colitis by promoting NLRP6-mediated autophagy. Cell Reports. 2022;41(2).

51. Li R, Zan Y, Wang D, Chen X, Wang A, Tan H, et al. A mouse model to distinguish NLRP6-mediated inflammasome-dependent and -independent functions. Proc Natl Acad Sci U S A. 2024;121(6):e2321419121.

52. Wang D, Chen X, Sui K, Wang A, Li R, Zhou W, et al. UBE2O-mediated monoubiquitination licenses NLRP6 inflammasome activation in the intestine. Cell Host Microbe. 2026;34(1):86–102 e8.

53. Anand PK, Malireddi RK, Lukens JR, Vogel P, Bertin J, Lamkanfi M, et al. NLRP6 negatively regulates innate immunity and host defence against bacterial pathogens. Nature. 2012;488(7411):389–93.

54. Seregin SS, Golovchenko N, Schaf B, Chen J, Eaton KA, Chen GY. NLRP6 function in inflammatory monocytes reduces susceptibility to chemically induced intestinal injury. Mucosal Immunol. 2017;10(2):434–45.

55. Lund ME, To J, O’Brien BA, Donnelly S. The choice of phorbol 12-myristate 13-acetate differentiation protocol influences the response of THP-1 macrophages to a pro-inflammatory stimulus. J Immunol Methods. 2016;430:64–70.

56. Genin M, Clement F, Fattaccioli A, Raes M, Michiels C. M1 and M2 macrophages derived from THP-1 cells differentially modulate the response of cancer cells to etoposide. BMC Cancer. 2015;15:577.

57. Ying W, Cheruku PS, Bazer FW, Safe SH, Zhou B. Investigation of macrophage polarization using bone marrow derived macrophages. J Vis Exp. 2013(76).

58. Vaudaux P, Waldvogel FA. Gentamicin antibacterial activity in the presence of human polymorphonuclear leukocytes. Antimicrob Agents Chemother. 1979;16(6):743–9.

59. Honma K, Mishima E, Inagaki S, Sharma A. The OxyR homologue in Tannerella forsythia regulates expression of oxidative stress responses and biofilm formation. Microbiology. 2009;155(Pt 6):1912–22.

60. Schindelin J, Arganda-Carreras I, Frise E, Kaynig V, Longair M, Pietzsch T, et al. Fiji: an open-source platform for biological-image analysis. Nat Methods. 2012;9(7):676–82.

61. Fernandez R, Moisy C. Fijiyama: a registration tool for 3D multimodal time-lapse imaging. Bioinformatics. 2021;37(10):1482–4.

62. Wingett SW, Andrews S. FastQ Screen: A tool for multi-genome mapping and quality control. F1000Res. 2018;7:1338.

63. Ewels P, Magnusson M, Lundin S, Kaller M. MultiQC: summarize analysis results for multiple tools and samples in a single report. Bioinformatics. 2016;32(19):3047–8.

64. Kim D, Langmead B, Salzberg SL. HISAT: a fast spliced aligner with low memory requirements. Nat Methods. 2015;12(4):357–60.

65. Liao Y, Smyth GK, Shi W. featureCounts: an efficient general purpose program for assigning sequence reads to genomic features. Bioinformatics. 2014;30(7):923–30.

66. Love MI, Huber W, Anders S. Moderated estimation of fold change and dispersion for RNA-seq data with DESeq2. Genome Biol. 2014;15(12):550.

67. Zhu A, Ibrahim JG, Love MI. Heavy-tailed prior distributions for sequence count data: removing the noise and preserving large differences. Bioinformatics. 2019;35(12):2084–92.

68. Wu T, Hu E, Xu S, Chen M, Guo P, Dai Z, et al. clusterProfiler 4.0: A universal enrichment tool for interpreting omics data. Innovation (Camb). 2021;2(3):100141.

69. Wilson E, Jackson S, Cruwys S, Kerry P. An evaluation of the immunohistochemistry benefits of boric acid antigen retrieval on rat decalcified joint tissues. J Immunol Methods. 2007;322(1-2):137–42.

70. Periasamy S, Kolenbrander PE. Mutualistic biofilm communities develop with Porphyromonas gingivalis and initial, early, and late colonizers of enamel. J Bacteriol. 2009;191(22):6804–11.

71. Lamont RJ, El-Sabaeny A, Park Y, Cook GS, Costerton JW, Demuth DR. Role of the Streptococcus gordonii SspB protein in the development of Porphyromonas gingivalis biofilms on streptococcal substrates. Microbiology. 2002;148(Pt 6):1627–36.

72. Yu S, Ding L, Liang D, Luo L. Porphyromonas gingivalis inhibits M2 activation of macrophages by suppressing alpha-ketoglutarate production in mice. Mol Oral Microbiol. 2018;33(5):388–95.

73. Lin J, Huang D, Xu H, Zhan F, Tan X. Macrophages: A communication network linking Porphyromonas gingivalis infection and associated systemic diseases. Front Immunol. 2022;13:952040.

74. Fleetwood AJ, Lee MKS, Singleton W, Achuthan A, Lee MC, O’Brien-Simpson NM, et al. Metabolic Remodeling, Inflammasome Activation, and Pyroptosis in Macrophages Stimulated by Porphyromonas gingivalis and Its Outer Membrane Vesicles. Front Cell Infect Microbiol. 2017;7:351.

75. Hajishengallis G, Martin M, Sojar HT, Sharma A, Schifferle RE, DeNardin E, et al. Dependence of bacterial protein adhesins on toll-like receptors for proinflammatory cytokine induction. Clin Diagn Lab Immunol. 2002;9(2):403–11.

76. Canton J, Khezri R, Glogauer M, Grinstein S. Contrasting phagosome pH regulation and maturation in human M1 and M2 macrophages. Mol Biol Cell. 2014;25(21):3330–41.

77. Gorska R, Gregorek H, Kowalski J, Laskus-Perendyk A, Syczewska M, Madalinski K. Relationship between clinical parameters and cytokine profiles in inflamed gingival tissue and serum samples from patients with chronic periodontitis. J Clin Periodontol. 2003;30(12):1046–52.

78. Zhou LN, Bi CS, Gao LN, An Y, Chen F, Chen FM. Macrophage polarization in human gingival tissue in response to periodontal disease. Oral Dis. 2019;25(1):265–73.

79. van Steenbergen TJ, Kastelein P, Touw JJ, de Graaff J. Virulence of black-pigmented Bacteroides strains from periodontal pockets and other sites in experimentally induced skin lesions in mice. J Periodontal Res. 1982;17(1):41–9.

80. Neiders ME, Chen PB, Suido H, Reynolds HS, Zambon JJ, Shlossman M, et al. Heterogeneity of virulence among strains of Bacteroides gingivalis. J Periodontal Res. 1989;24(3):192–8.

81. Hasturk H, Kantarci A, Van Dyke TE. Oral inflammatory diseases and systemic inflammation: role of the macrophage. Front Immunol. 2012;3:118.

82. Graves D. Cytokines that promote periodontal tissue destruction. J Periodontol. 2008;79(8 Suppl):1585–91.

83. Yu G, Choi YK, Lee S. Inflammasome diversity: exploring novel frontiers in the innate immune response. Trends Immunol. 2024.

84. Sundaram B, Tweedell RE, Prasanth Kumar S, Kanneganti TD. The NLR family of innate immune and cell death sensors. Immunity. 2024;57(4):674–99.

85. Didilescu AC, Chinthamani S, Scannapieco FA, Sharma A. NLRP3 inflammasome activity and periodontal disease pathogenesis-A bidirectional relationship. Oral Dis. 2024;30(7):4069–77.

86. Zhang J, Liu X, Wan C, Liu Y, Wang Y, Meng C, et al. NLRP3 inflammasome mediates M1 macrophage polarization and IL-1beta production in inflammatory root resorption. J Clin Periodontol. 2020.

87. Rocha FRG, Delitto AE, de Souza JAC, Gonzalez-Maldonado LA, Wallet SM, Rossa Junior C. Relevance of Caspase-1 and Nlrp3 Inflammasome on Inflammatory Bone Resorption in A Murine Model of Periodontitis. Sci Rep. 2020;10(1):7823.

88. Bostanci N, Meier A, Guggenheim B, Belibasakis GN. Regulation of NLRP3 and AIM2 inflammasome gene expression levels in gingival fibroblasts by oral biofilms. Cell Immunol. 2011;270(1):88–93.

89. Yamaguchi Y, Kurita-Ochiai T, Kobayashi R, Suzuki T, Ando T. Regulation of the NLRP3 inflammasome in Porphyromonas gingivalis-accelerated periodontal disease. Inflamm Res. 2017;66(1):59–65.

90. Chen Y, Yang Q, Lv C, Chen Y, Zhao W, Li W, et al. NLRP3 regulates alveolar bone loss in ligature-induced periodontitis by promoting osteoclastic differentiation. Cell Prolif. 2021;54(2):e12973.

91. Okano T, Ashida H, Suzuki S, Shoji M, Nakayama K, Suzuki T. Porphyromonas gingivalis triggers NLRP3-mediated inflammasome activation in macrophages in a bacterial gingipains-independent manner. Eur J Immunol. 2018;48(12):1965–74.

92. Aral K, Milward MR, Kapila Y, Berdeli A, Cooper PR. Inflammasomes and their regulation in periodontal disease: A review. J Periodontal Res. 2020;55(4):473–87.

93. Mark Welch JL, Rossetti BJ, Rieken CW, Dewhirst FE, Borisy GG. Biogeography of a human oral microbiome at the micron scale. Proc Natl Acad Sci U S A. 2016;113(6):E791–800.

94. Costalonga M, Herzberg MC. The oral microbiome and the immunobiology of periodontal disease and caries. Immunol Lett. 2014;162(2 Pt A):22–38.

95. Nowicki EM, Shroff R, Singleton JA, Renaud DE, Wallace D, Drury J, et al. Microbiota and Metatranscriptome Changes Accompanying the Onset of Gingivitis. MBio. 2018;9(2).

96. Griffen AL, Beall CJ, Campbell JH, Firestone ND, Kumar PS, Yang ZK, et al. Distinct and complex bacterial profiles in human periodontitis and health revealed by 16S pyrosequencing. ISME J. 2012;6(6):1176–85.

97. Hajishengallis G, Lamont RJ. Polymicrobial communities in periodontal disease: Their quasi-organismal nature and dialogue with the host. Periodontol 2000. 2021;86(1):210–30.

98. Varanat M, Haase EM, Kay JG, Scannapieco FA. Activation of the TREM-1 pathway in human monocytes by periodontal pathogens and oral commensal bacteria. Mol Oral Microbiol. 2017;32(4):275–87.

99. Bynum KT, Panasiewicz M, Kay JG. Neutrophil Activation Decreases Ability to Kill Oral Streptococcus gordonii. Mol Oral Microbiol. 2026;41(2):94–106.

100. Pelegrin P, Barroso-Gutierrez C, Surprenant A. P2X7 receptor differentially couples to distinct release pathways for IL-1beta in mouse macrophage. J Immunol. 2008;180(11):7147–57.

101. Cecil JD, O’Brien-Simpson NM, Lenzo JC, Holden JA, Singleton W, Perez-Gonzalez A, et al. Outer Membrane Vesicles Prime and Activate Macrophage Inflammasomes and Cytokine Secretion In Vitro and In Vivo. Front Immunol. 2017;8:1017.

102. Almeida-da-Silva CLC, Ramos-Junior ES, Morandini AC, Rocha GDC, Marinho Y, Tamura AS, et al. P2X7 receptor-mediated leukocyte recruitment and Porphyromonas gingivalis clearance requires IL-1beta production and autocrine IL-1 receptor activation. Immunobiology. 2018:1–0.

103. Choi CH, Spooner R, DeGuzman J, Koutouzis T, Ojcius DM, Yilmaz O. Porphyromonas gingivalis-nucleoside-diphosphate-kinase inhibits ATP-induced reactive-oxygen-species via P2X7 receptor/NADPH-oxidase signalling and contributes to persistence. Cell Microbiol. 2013;15(6):961–76.

104. Ramos-Junior ES, Morandini AC, Almeida-da-Silva CL, Franco EJ, Potempa J, Nguyen KA, et al. A Dual Role for P2X7 Receptor during Porphyromonas gingivalis Infection. J Dent Res. 2015;94(9):1233–42.

105. Park E, Na HS, Song YR, Shin SY, Kim YM, Chung J. Activation of NLRP3 and AIM2 inflammasomes by Porphyromonas gingivalis infection. Infect Immun. 2014;82(1):112–23.

106. Huang CB, Alimova Y, Ebersole JL. Macrophage polarization in response to oral commensals and pathogens. Pathog Dis. 2016;74(3):ftw011.

107. Huang CB, Altimova Y, Strange S, Ebersole JL. Polybacterial challenge effects on cytokine/chemokine production by macrophages and dendritic cells. Inflamm Res. 2011;60(2):119–25.

108. Abe T, Hajishengallis G. Optimization of the ligature-induced periodontitis model in mice. J Immunol Methods. 2013;394(1-2):49–54.

109. Graves DT, Fine D, Teng YT, Van Dyke TE, Hajishengallis G. The use of rodent models to investigate host-bacteria interactions related to periodontal diseases. J Clin Periodontol. 2008;35(2):89–105.

110. Duarte C, Yamada C, Ngala B, Garcia C, Akkaoui J, Birsa M, et al. Effects of IL-34 and anti-IL-34 neutralizing mAb on alveolar bone loss in a ligature-induced model of periodontitis. Mol Oral Microbiol. 2023.

111. Tan J, Patil PC, Luzzio FA, Demuth DR. In Vitro and In Vivo Activity of Peptidomimetic Compounds That Target the Periodontal Pathogen Porphyromonas gingivalis. Antimicrob Agents Chemother. 2018;62(7).

112. Sohn J, Li L, Zhang L, Settem RP, Honma K, Sharma A, et al. Porphyromonas gingivalis indirectly elicits intestinal inflammation by altering the gut microbiota and disrupting epithelial barrier function through IL9-producing CD4(+) T cells. Mol Oral Microbiol. 2022;37(2):42–52.

113. Xiao W, Dong G, Pacios S, Alnammary M, Barger LA, Wang Y, et al. FOXO1 deletion reduces dendritic cell function and enhances susceptibility to periodontitis. Am J Pathol. 2015;185(4):1085–93.

114. Wlodarska M, Thaiss CA, Nowarski R, Henao-Mejia J, Zhang JP, Brown EM, et al. NLRP6 inflammasome orchestrates the colonic host-microbial interface by regulating goblet cell mucus secretion. Cell. 2014;156(5):1045–59.

115. Ghimire L, Paudel S, Jin L, Baral P, Cai S, Jeyaseelan S. NLRP6 negatively regulates pulmonary host defense in Gram-positive bacterial infection through modulating neutrophil recruitment and function. PLoS Pathog. 2018;14(9):e1007308.

116. Lamont RJ, Hajishengallis G. Polymicrobial synergy and dysbiosis in inflammatory disease. Trends Mol Med. 2015;21(3):172–83.

117. Mark Welch JL, Dewhirst FE, Borisy GG. Biogeography of the Oral Microbiome: The Site-Specialist Hypothesis. Annu Rev Microbiol. 2019;73:335–58.

118. Elinav E, Strowig T, Kau AL, Henao-Mejia J, Thaiss CA, Booth CJ, et al. NLRP6 inflammasome regulates colonic microbial ecology and risk for colitis. Cell. 2011;145(5):745–57.

119. Ghimire L, Paudel S, Jin L, Jeyaseelan S. The NLRP6 inflammasome in health and disease. Mucosal Immunol. 2020;13(3):388–98.

120. Liu W, Liu J, Wang W, Wang Y, Ouyang X. NLRP6 Induces Pyroptosis by Activation of Caspase-1 in Gingival Fibroblasts. J Dent Res. 2018;97(12):1391–8.

121. Lu WL, Zhang L, Song DZ, Yi XW, Xu WZ, Ye L, et al. NLRP6 suppresses the inflammatory response of human periodontal ligament cells by inhibiting NF-kappaB and ERK signal pathways. Int Endod J. 2019;52(7):999–1009.

122. Khoury W, Glogauer J, Tenenbaum HC, Glogauer M. Oral inflammatory load: Neutrophils as oral health biomarkers. J Periodontal Res. 2020;55(5):594–601.

123. Elebyary O, Sun C, Batistella EA, Van Dyke TE, Low SB, Singhal S, et al. Utilizing Oral Neutrophil Counts as an Indicator of Oral Inflammation Associated With Periodontal Disease: A Blinded Multicentre Study. J Clin Periodontol. 2024;51(11):1410–20.

124. Moutsopoulos NM, Chalmers NI, Barb JJ, Abusleme L, Greenwell-Wild T, Dutzan N, et al. Subgingival microbial communities in Leukocyte Adhesion Deficiency and their relationship with local immunopathology. PLoS Pathog. 2015;11(3):e1004698.

125. Dutzan N, Kajikawa T, Abusleme L, Greenwell-Wild T, Zuazo CE, Ikeuchi T, et al. A dysbiotic microbiome triggers T(H)17 cells to mediate oral mucosal immunopathology in mice and humans. Sci Transl Med. 2018;10(463).

126. Christ A, Gunther P, Lauterbach MAR, Duewell P, Biswas D, Pelka K, et al. Western Diet Triggers NLRP3-Dependent Innate Immune Reprogramming. Cell. 2018;172(1-2):162–75 e14.

127. Hajishengallis G. Epigenetic inflammatory memory and periodontal disease: Mechanisms and clinical significance for comorbidities. J Periodontol. 2025.

