## Supplemental Figure 1 for "Polymicrobial-driven NLRP6 inflammasome regulates IL-1β production and alveolar bone loss in a murine model of periodontitis"

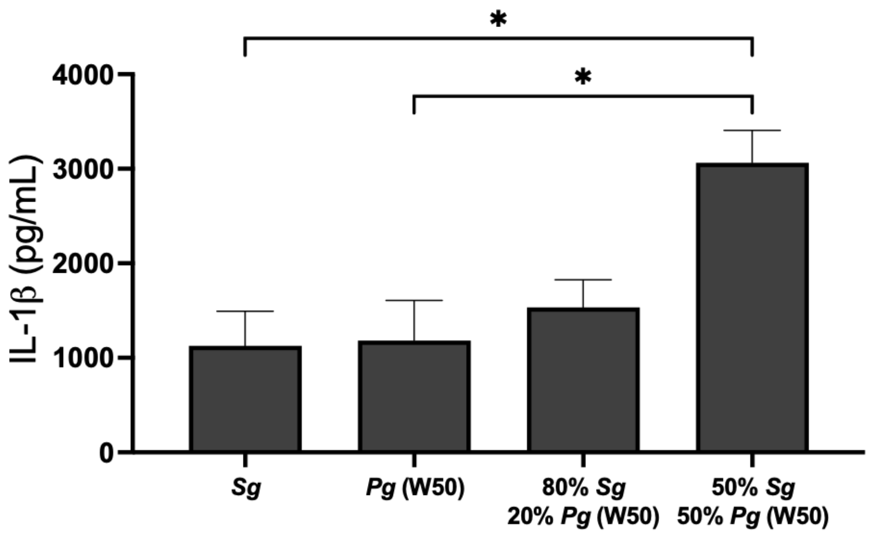


**Figure S1. IL-1β release from macrophages increased with co-incubation of P. gingivalis strain W50.** IL-1β release from PMA differentiated THP-1 cells incubated with S. gordonii, P. gingivalis strain W50, or a combination thereof (all total MOI = 10:1) after 6 hours incubation. Shown are means ±SEM of 3 independent experiments. P values (*<0.05) were calculated by ordinary one-way ANOVA followed by Holm-Šídák's multiple comparisons tests.
